# Evidence of recombination in coronaviruses implicating pangolin origins of nCoV-2019

**DOI:** 10.1101/2020.02.07.939207

**Authors:** Matthew C. Wong, Sara J. Javornik Cregeen, Nadim J. Ajami, Joseph F. Petrosino

## Abstract

A novel coronavirus (nCoV-2019) was the cause of an outbreak of respiratory illness detected in Wuhan, Hubei Province, China in December of 2019. Genomic analyses of nCoV-2019 determined a 96% resemblance with a coronavirus isolated from a bat in 2013 (RaTG13); however, the receptor binding motif (RBM) of these two genomes share low sequence similarity. This divergence suggests a possible alternative source for the RBM coding sequence in nCoV-2019. We identified high sequence similarity in the RBM between nCoV-2019 and a coronavirus genome reconstructed from a viral metagenomic dataset from pangolins possibly indicating a more complex origin for nCoV-2019.

---

An outbreak of respiratory illness caused by a novel coronavirus (nCoV-2019)^1^, first identified in Wuhan China, has resulted in over twenty thousand confirmed cases worldwide (WHO Situation Report, February 5 2020). Data available to date has led to the hypothesis that the outbreak strain originated from bats. The nCoV-2019 has been reported to share 96% sequence identity to the RaTG13 genome, a coronavirus isolated from an intermediate horseshoe bat (*Rhinolophus affinis*)^2^. RaTG13 is the most closely related coronavirus genome to nCoV-2019, however other viral strains may have contributed to its genomic content. To explore this hypothesis, we examined several viral metagenomic datasets for potential coronaviruses that could harbor RBMs closer to nCoV-2019. We report evidence of a coronavirus recovered from a Malayan pangolin (*Manis javanica*) viral metagenomic dataset^3^ that shares a higher sequence identity at a crucial segment of the genome involved in host infection (Figure 1). This result indicates a potential recombination event and a more complex origin for the nCoV-2019 strain.

**Figure 1:**
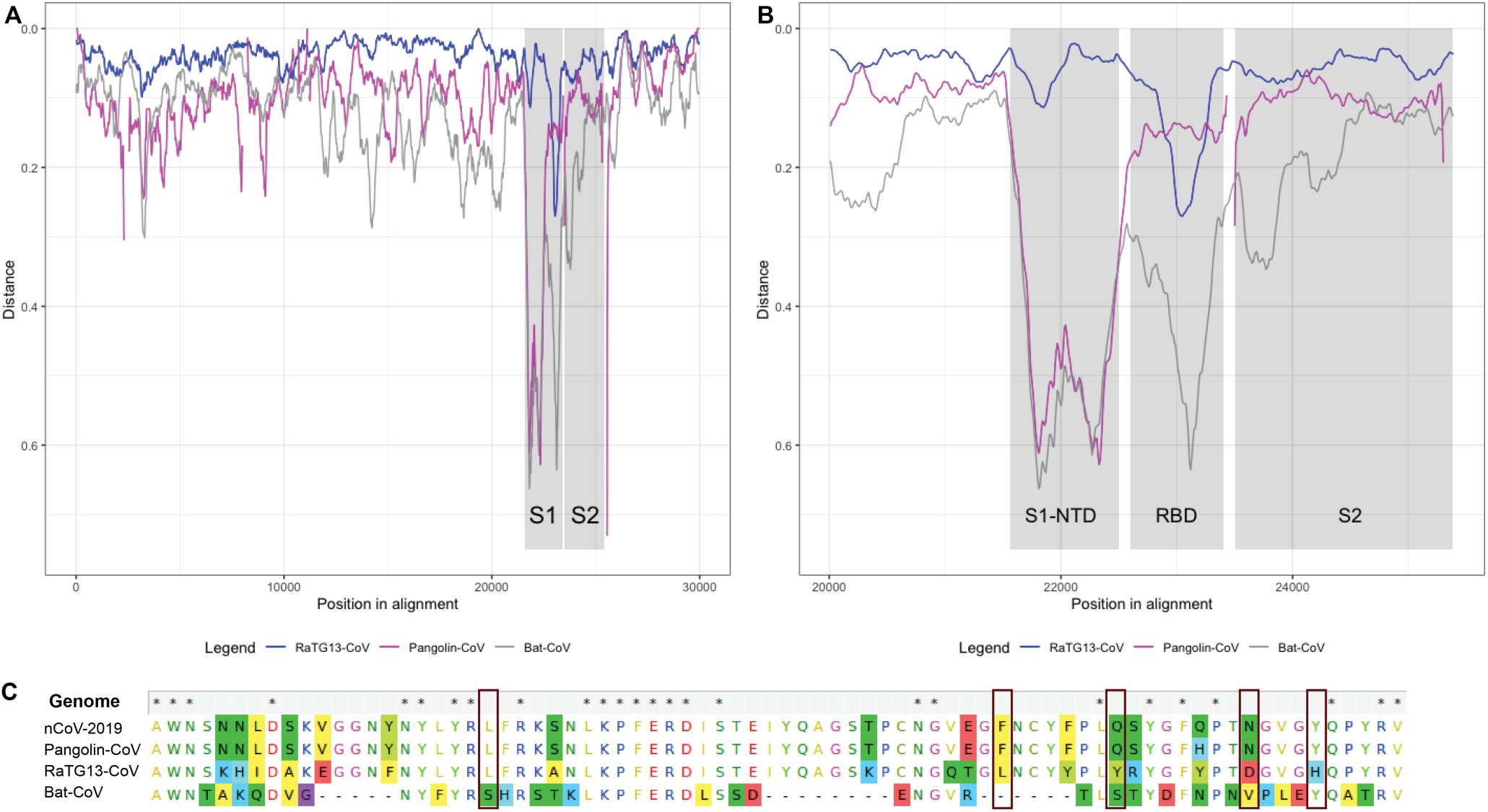
Nucleotide and amino acid identity across three coronavirus genomes relative to the outbreak strain (nCoV-2019). Panel A: Distances of selected coronavirus genomes relative to nCov-2019 across the entire genome length. The gaps in the Pangolin-CoV are a result of the incompleteness of the genome used for comparison. Grey box indicates S1 and S2 domains of the spike protein. Panel B: Close up of the S1 domain with surrounding regions. Grey boxes highlight the S1-NTD subdomain and the S1-CTD or RBD subdomain. In the RBD subdomain the homology of the Pangolin-CoV genome exceeds that of RaTG13-CoV as compared to the nCoV-2019 strain. Panel C: Multiple alignment of the receptor biding motif at the amino acid level. Boxes highlight the key amino acid residues involved in ACE2 receptor binding. The nCoV-2019 and Pangolin-CoV strains are identical at all five amino acid residues, whereas RaTG13-CoV strain differs at four out of five compared to nCoV-2019. Labels: nCoV-2019 – outbreak strain; Pangolin-CoV – pangolin coronavirus identified in this study; RaTG13-CoV – *R. affinis* coronavirus; Bat-CoV – bat coronavirus (MG772933.1)

The coronavirus spike protein is a trimeric protein that mediates virus entry into the host cell. It is composed of two domains: S1 and S2. The S1 domain contains two subdomains: S1-NTD and S1-CTD, also known as the receptor binding domain (RBD). The S2 domain encodes the stalk of the spike protein and is highly conserved across the SARS-like coronaviruses^4^. In order to infect a host cell, the RBD must first bind to a surface protein on the host cell. In the case of SARS-CoV and nCoV-2019, this is the angiotensin converting enzyme 2 (ACE2) receptor, expressed in human airway epithelia as well as lung parenchyma among other tissues. Following RBD binding, the S1/S2 junction must be cleaved by a surface protease^4^. Zhou et al. (2020)^2^ have reported noticeable decreased amino acid homology between the RBD of the spike protein in RaTG13 and nCoV-2019. A further analysis of the RBD in RaTG13 showed the majority of the divergence was restricted to amino acid residues 435 to 510 (75% nucleotide identity and 78% amino acid identity). This region corresponds to the receptor binding motif (RBM), which is responsible for host specificity. Such divergence at the RBM could indicate the possibility of an alternate source for the RBM coding sequence in nCoV-2019. To explore this, we examined several viral metagenomic datasets for potential coronaviruses that could harbor RBMs closer to nCoV-2019. Using VirMAP^5^, we aimed to reconstruct coronavirus genomes to compare recovered RBM sequences to that of nCoV-2019.

## Methods

### Data used in the study

Viral metagenomic datasets of hosts that could potentially harbor coronaviruses were downloaded from the NCBI BioProject database. These included PRJNA573298^3^ (pangolin), PRJNA597258 (fruit bats) and PRJNA379515 (bats). The coronavirus genome (recovered from bat, GenBank accession number MG772933.1^6^) reported to be the most homologous to the nCoV-2019 outbreak strain prior to the publication the RaTG13 (GenBank accession number MN996532) was also downloaded.

### Analysis

Sequences from each project were trimmed (minimum quality score 20 and minimum length 75bp) and filtered for Illumina adapter and PhiX presence using bbduk.sh v38.71^7^. Prior to analysis all sequences were filtered against a the relevant host reference genome (pangolin ManJav1.0, PRJNA256023 or bat mRhiFer1_v1.p, PRJNA489106) using bbmap.sh v38.71(k=15 and minid=0.9)^7^. After trimming and filtering, sequence reads were processed using VirMAP^5^ to obtain and identify coronavirus genomes present in the viral metagenomic dataset. Two coronavirus genomes sharing high homology to nCoV-2019 were recovered from the pangolin dataset in two samples (SRR10168377 and SRR10168378). The two genomes were merged using the easymerge.pl subcommand from VirMAP to create the final pangolin-associated coronavirus (Pangolin-CoV) genome.

All further analysis was done using four coronaviruses genome sequences: Pangolin-CoV, RaTG13, Bat-CoV and nCoV-2019. They were pairwise aligned on NCBI using the blastn and blastx algorithms^8,9^ and sequence homology plots were calculated using RDP4 beta 99^10^. Distances of the selected coronavirus genomes relative to nCov-2019 were calculated using a 300bp window sliding every 10bp across the entire genome length. Sequence homology plots were visualized using R version 3.6.2^11^ and the ggplot2 visualization package^12^. The RBM segments as well as the Pro-Arg-Arg-Ala (PRRA) amino acid insertion region were excised from the full genome and pairwise aligned on NCBI using blastn and blastx. Multiple alignments of the both these regions were done using MAFFT v7.313^13^ and visualized with MEGAX v10.1.7^14^.

## Results

Of the three viral metagenomic datasets inspected, only the pangolin dataset contained coronavirus genomes sharing significant homology with the outbreak strain. Two samples in this dataset yielded two partial coronavirus genomes that overlapped >8.4kb at >99% nucleotide identity with each other. After merging, the final Pangolin-CoV draft genome was 90.5% similar to nCoV-2019 (median nucleotide identity). Figure 1 shows the sequence homology of Pangolin-CoV, RaTG13-CoV and Bat-CoV relative to nCoV-2019 (Figure 1A shows the homology across the entire genome length). The multiple alignments across the RBM segments revealed an 89% nucleotide (Figure 1B) and 98% amino acid identity (Figure 1C) of the Pangolin-CoV compared to nCoV-2019. The sequence homology plots showed that the Pangolin-CoV was more similar to nCoV-2019 at the RBD region than the RaTG13-CoV and Bat-CoV (Figure 1B). The multiple alignment comparing the PRRA insertion region at the S1/S2 junction shows the insertion present in nCoV-2019 and missing in the other three CoV genomes (Figure 2).

**Figure 2:**
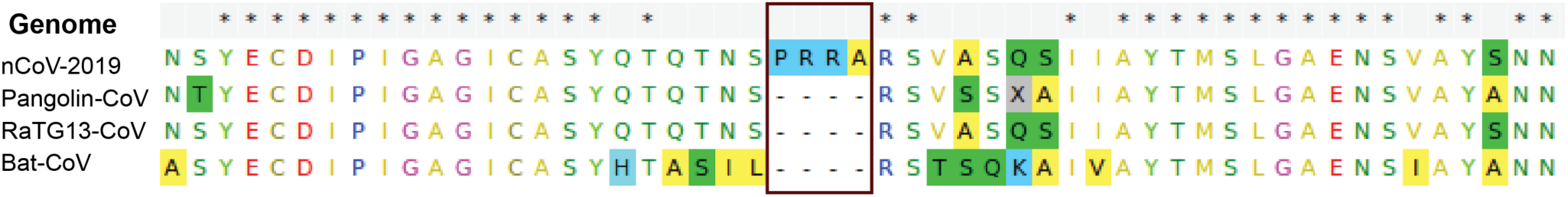
Multiple alignment at the amino acid level of PRRA insertion region and flanking areas across three coronavirus genomes relative to the outbreak strain (nCoV-2019). Box show the PRRA insertion present in nCoV-2019 and missing in the other three CoV genomes.

## Discussion

Recent coronavirus zoonotic transmissions have involved non-bat intermediates. Indeed, both SARS^15^ and MERS^16^ are believed to have arisen from coronaviruses recombining in civet cats and camels, respectively. While early indications for nCoV-2019 showed that the virus may have jumped directly from bats to humans^2^, here we present evidence that nCoV-2019 likely arose from yet another recombined coronavirus in an intermediate host. A dataset describing the viral diversity in Malayan pangolins^3^ yielded a coronavirus genome that shares 89% nucleotide (Figure 1A and 1B) and 98% amino acid (Figure 1C) identity across the same RBM segment with nCoV-2019. Additionally, the RaTG13 genome only shares one out of five key amino acids involved in receptor binding motif^2^, whereas the pangolin coronavirus shares all five key residues (Figure 1C). It therefore appears that the RBM in nCoV-2019 was introduced during a recombination event between the strain of coronavirus found in the pangolin and RaTG13. Given the heavy selective pressure under which the RBM is constrained, it is unlikely that RaTG13 acquired the mutations found in the pangolin coronavirus genome through random chance. We posit that the homology at the RBM between nCoV-2019 and a pangolin coronavirus may have arisen from a recombination event based on sequence analysis presented herein.

Amino acid alignment between pangolin coronavirus and nCoV-2019 at the S1-NTD subdomain results in low sequence similarity. However, increased similarity is observed from the RBD through the S2 domain (Figure 1A-B). This suggests a possible recombination event between a bat coronavirus and the pangolin coronavirus at the S1-NTD and RBD junction. While there is high conservation at the amino-acid level between the pangolin coronavirus and nCoV-2019, this is not reflected in the nucleotide identity, suggesting that the proposed recombination event occurred in the more distant past, allowing subsequent time for genetic drift. One such mutation could be the insertion of the amino acid residues PRRA near the S1/S2 junction which induces a furin cleavage motif. The secondary requirement for entry into the host cell is proteolytic cleavage of the S1/S2 domains in order to mediate membrane fusion post ACE2 binding ^4^, and introduction of the furin cleavage motif may be another factor in nCoV-2019 being able to replicate in humans (Figure 2).

The main limitations of this analysis are the lack of additional viral datasets arising from pangolins or the availability of coronavirus genomes isolated from pangolins that could be used for multiple genome alignments with nCoV-2019. In addition, the current analysis only focuses on the spike protein of nCoV-2019 without exploring additional recombination events that may lead to changes in host tropism. Nonetheless, the identification of a possible recombination event in the host-receptor binding protein of nCoV-2019, suggesting an intermediate host, supports a call for continuous surveillance of species known to be infected by this group of viruses, including pangolins, as well as all small mammals sold in such wet markets as the Huanan Seafood Wholesale Market in Wuhan to better understand and characterize zoonotic transmission events.

## Acknowledgements

Research reported in this publication was supported by the National Institute Of Allergy And Infectious Diseases of the National Institutes of Health under Award Number U19AI144297. The authors thank the Alkek foundation for ongoing support of the Baylor College of Medicine Alkek Center for Metagenomics and Microbiome Research. We thank P. Hotez, R. Atmar, and G. Metcalf for critical review of the manuscript.

## References

1. Wu F, Zhao S, Yu B, Chen YM, Wang W, Song ZG, Hu Y et al. A new coronavirus associated with human respiratory disease in China. Nature 2020;453

2. Zhou P, Yang XL, Wang XG, Hu B, Zhang L, Zhang W et al. A pneumonia outbreak associated with a new coronavirus of probable bat origin. Nature 2020;1221

3. Liu P, Chen W, Chen JP. Viral Metagenomics Revealed Sendai Virus and Coronavirus Infection of Malayan Pangolins (*Manis javanica*). Viruses 2019; 11(11): 979

4. Li, F. Structure, Function, and Evolution of Coronavirus Spike Proteins. Annual Review of Virology 2016;3(1): 237–261

5. Ajami NJ, Wong MC, Ross MC, Lloyd RE, Petrosino JF. Maximal viral information recovery from sequence data using VirMAP. Nature Communications 2018; 9: 3205

6. Hu D, Zhu C, Ai L, He T, Wang Y, Ye F, et al. Genomic characterization and infectivity of a novel SARS-like coronavirus in Chinese bats. Emerging microbes and infections. 2018; 7(1): 1–10

7. Bushnell B. BBMap Short Read Aligner. Berkeley, California: University of California; 2016; sourceforge.net/projects/bbmap/

8. Altschul SF, Gish W, Miller W, Myers EW, Lipman DJ. Basic local alignment search tool. Journal of Molecular Biology 1990; 215:403–410.

9. Camacho C, Coulouris G, Avagyan V, Ma N, Papadopoulos J, Bealer K, Madden TL. BLAST+: architecture and applications. BMC Bioinformatics 2008; 10:421.

10. Martin DP, Murrell B, Golden M, Khoosal A, Muhire B. RDP4: Detection and analysis of recombination patterns in virus genomes. Virus Evolution 2015; 1–5

11. R Core Team (2019). R: A language and environment for statistical computing. R Foundation for Statistical Computing, Vienna, Austria. URL https://www.R-project.org/

12. H. Wickham. ggplot2: Elegant Graphics for Data Analysis. Springer-Verlag New York, 2016.

13. Katoh K, Standley DM. MAFFT Multiple Sequence Alignment Software Version 7: Improvements in Performance and Usability. Molecular Biology and Evolution 2013; 30(4): 772–780.

14. Kumar S, Stecher G, Li M, Knyaz C, Tamura K. MEGA X: Molecular Evolutionary Genetics Analysis across computing platforms. Molecular Biology and Evolution 2018; 35:1547–1549.

15. Lau SK, Feng Y, Chen H, et al. Severe Acute Respiratory Syndrome (SARS) Coronavirus ORF8 Protein Is Acquired from SARS-Related Coronavirus from Greater Horseshoe Bats through Recombination. Journal of Virology 2015;89(20):10532–10547.

16. Zhang Z, Shen L, Gu X. Evolutionary Dynamics of MERS-CoV: Potential Recombination, Positive Selection and Transmission. Scientific Reports 2016; 6:25049.

